# Citrullination regulates wound responses and tissue regeneration in zebrafish

**DOI:** 10.1101/2019.12.27.889378

**Authors:** Netta Golenberg, Jayne M. Squirrell, David A. Bennin, Julie Rindy, Paige E. Pistono, Kevin W. Eliceiri, Miriam A. Shelef, Junsu Kang, Anna Huttenlocher

## Abstract

Calcium signaling is an important early step in wound healing, yet how these early signals promote regeneration remains unclear. Peptidylarginine deiminases (PADs), a family of calcium-dependent enzymes, catalyze citrullination, a post-translational modification that alters protein function and has been implicated in autoimmune diseases. We generated a mutation in the single zebrafish ancestral *pad* gene, *padi2,* resulting in a loss of detectable calcium-dependent citrullination. The *padi2* mutants exhibit impaired resolution of inflammation and regeneration after caudal fin transection. Further, we identified a new subpopulation of cells displaying citrullinated histones within the notochord bead following tissue injury. Citrullination of histones in this region was absent and wound-induced proliferation was perturbed in Padi2-deficient larvae. Taken together, our results show that Padi2 is required for the citrullination of histones within a group of cells in the notochord bead, and for promoting wound-induced proliferation required for efficient regeneration. These findings identify Padi2 as a potential intermediary between early calcium signaling and subsequent tissue regeneration.

**Summary:** Golenberg et al. developed a citrullination-deficient zebrafish and demonstrated a role for Padi2 in fin wound responses and regeneration. This work identified a distinct population of cells within the regenerative notochord bead that exhibited wound-induced histone citrullination.

## Introduction

Humans have limited regenerative capacity, resulting in injury-induced scarring and loss of tissue function in response to damage that can present significant clinical challenges. Because mammalian wound repair occurs through similar stages as regeneration in simple animal models (Yokoyama, 2008), regenerative animal models may provide insight into the molecular signals that optimize mammalian wound healing. After wounding, early signals activate a series of regenerative steps. The initial wound-healing phase is defined by the migration and formation of the wound epithelium and the recruitment of immune cells (Roehl, 2018). This is followed by the regenerative program, including the formation of a blastema, a mass of stem-cell like cells that mediate cell proliferation and restoration of damaged tissue (Whitehead et al., 2005). Improper activation or regulation of these tightly controlled steps results in improper regeneration. Although increased cytosolic calcium early in wound healing has been linked to later regenerative proliferation (Globus et al., 1987; Lagoudakis et al., 2010; Yoo et al., 2012a), there is limited understanding of how these early signals impact later regenerative events.

An attractive candidate to link calcium increase with subsequent regeneration is the family of calcium-dependent enzymes, peptidylarginine deiminases (PADIs or PADs), which catalyze the deimination of a peptidyl-arginine to the neutrally-charged, non-coded amino acid, citrulline (Vossenaar et al., 2003). These enzymes were recently implicated in stem cell pluripotency because citrullination of histones and chromatin modifiers can maintain pluripotency by promoting an open chromatin state, thereby regulating gene expression (Christophorou et al., 2014; Wiese et al., 2019; Xiao et al., 2017). Increased expression and activity of PAD enzymes is associated with autoimmune disorders, cancers, and neurodegenerative disorders (Chang et al., 2009; Gyorgy et al., 2006). While PADs have been studied in mammalian models, the presence of multiple PAD isoforms and functional redundancies make it challenging to dissect the role of citrullination in normal development and tissue repair.

Zebrafish, *Danio rerio*, have one highly-conserved copy of a *pad* gene, *padi2*, that shares canonical mammalian PAD features, with conserved enzymatic activity and calcium dependence. We generated a *padi2* mutant zebrafish line lacking detectable calcium-dependent citrullination activity while displaying normal developmental but impaired regenerative growth. This work provides insight into how calcium-dependent citrullination may integrate early signals induced by injury to mediate subsequent tissue repair.

## Results and discussion

### Characterization of zebrafish peptidylarginine deiminase

To examine the role of citrullination in zebrafish, we first characterized the annotated zebrafish *pad* gene, *padi2*. A 7-exon transcript (203) and two 16-exon transcripts with alternative start sites (201 and 202) are annotated (Fig S1 A). The 7-exon transcript is predicted to lack the catalytic C-terminus, therefore, we focused on cloning the transcripts with the two predicted alternative start sites and the shared exon 16 and identified two splice variants of *padi2*. These transcripts have different first exons, but both splice variants share a complete exon 10, contrary to the genome assembly predictions (Fig S1 A, arrows). Due to these discrepancies, these transcripts are referred to here as 201a and 202 (Fig S1 A). These transcripts are highly conserved with human PADs and share 55% amino acid identity and 69% similarity to human PAD2 with conserved catalytic, substrate-binding and calcium-coordinating amino acids (Fig S1 B) (Smith and Waterman, 1981). Using our newly generated polyclonal antibody, immunoblotting showed a doublet at 75-80 kDa, providing further evidence for two full-length zebrafish *padi2* splice variants (Fig S1 C, arrow). Notably, this antibody did not detect a protein of equivalent size to the predicted transcript 203 (∼35 kDA) (Fig S1 C). The absence of this doublet with pre-immune serum and the detection of an appropriately sized protein in *padi2*-201a mRNA-injected larvae demonstrates the specificity of the antibody (Fig S1 D and E). This antibody also detected a large protein species at roughly 200 kDa that is of unclear significance (Fig S1 C, asterisk).

To assess citrullination activity, we used a colorimetric *in vitro* citrullination activity assay and detected citrullination activity with both bacterially expressed Padi2 variants (Fig 1 A) (Nakayama-Hamada et al., 2005). Mammalian PADs bind 5-6 calcium ions for proper structure and activity, with essential roles for Ca1 and Ca2 (Arita et al., 2004). Similarly, zebrafish Padi2 activity was also calcium dependent (Fig 1 A). Furthermore, alanine point mutations of amino acids predicted to be necessary for Ca1 and Ca2 binding sites (Arita et al., 2004) impaired *in vitro* citrullination activity (Fig 1 B and Fig S1 F). Mutation of the catalytic cysteine also abolished activity (Fig 1 B and Fig S1 F). These data indicate that zebrafish Padi2 is a canonical PAD with similar function as mammalian PAD enzymes.

**Figure 1:**
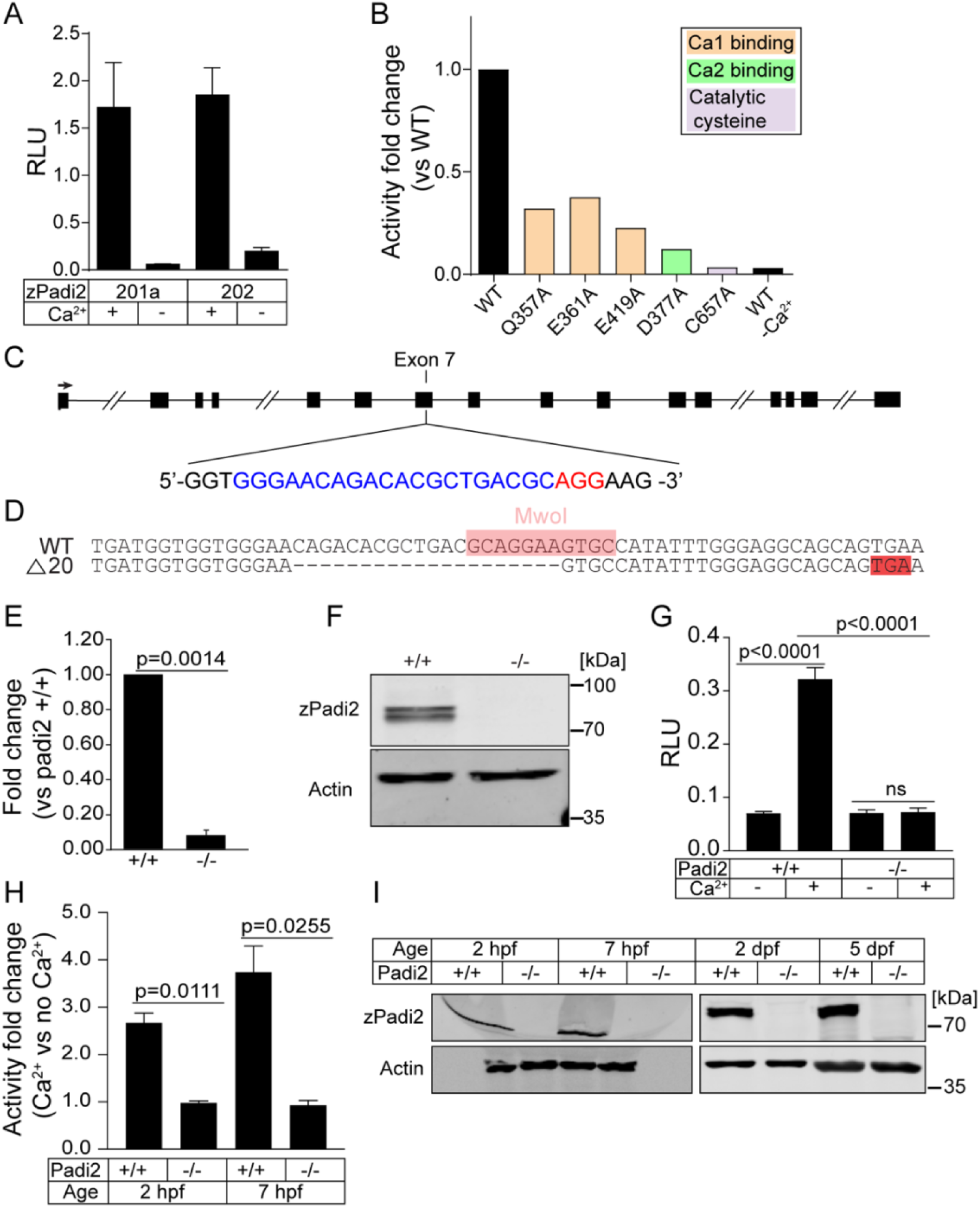
Characterization of zebrafish Padi2. (A) Citrullination activity of bacterially expressed zebrafish Padi2 201a and 202 splice variants in total lysates with and without calcium. Absorbance of light was measured and expressed as mean (± SEM) relative light units (RLU), normalized for protein level. Data represent 3 independent replicates. (B) Citrullination activity of Padi2 201a and individual point mutations in calcium binding and catalytic amino acids. Fold change of enzymatic activity is shown relative to wild-type Padi2 201a. Data represent 2 independent replicates and wild-type values are also represented in A. (C) Schematic of *padi2* gene with exon 7 gRNA sequence highlighted for CRISPR/Cas9 mutagenesis. gRNA sequence in blue, PAM site in red. (D) Sequence alignment of wild-type and *padi2*^−/−^ 20 bp mutation in exon 7. MwoI restriction site for genotyping highlighted in pink, predicted early stop codon highlighted in red. (E) RT-qPCR of *padi2* exon5/6 on individual larvae from a *padi2*^+/-^ incross. Data are from three pooled independent replicates with the means and SEM reported and a one-sample t test performed. (F) Representative western blot for zebrafish Padi2 and Actin from pooled larvae (representative of 4 experiments). (G) Citrullination activity of pooled zebrafish lysates expressed as relative light units (RLU). Data are from 3 independent replicates with the means and SEM reported and an ANOVA performed. (H) Citrullination activity of pooled embryo lysates during development. Fold change of enzymatic activity is shown as a ratio of calcium-treated to no calcium for each condition. Data are from 3 independent replicates. (I) Representative western blot for zebrafish Padi2 and Actin from pooled zebrafish through stages of development (representative of 3 and 2 experiments).

### Generation of a *padi2* zebrafish mutant

Padi2 is the ancestral protein of the mammalian PADs, with the broadest tissue distribution and substrate specificity in mammals (Rebl et al., 2010; Vossenaar et al., 2003). To characterize the role of citrullination in regeneration, we generated a zebrafish *padi* mutant using CRISPR/Cas9 gene editing, targeting exon 7 of *padi2*, an optimal target region preceding the catalytic amino acids and essential calcium binding sites. The resulting CRISPR-generated product had a 20 base pair deletion (Fig 1 C and D) and caused a predicted frameshift mutation resulting in an early stop codon. Padi2 homozygous mutants (*padi2*^−/−^) had reduced levels of *padi2* mRNA (Fig 1 E) and loss of Padi2 protein (Fig 1 F and Fig S1 C) at 2 days post fertilization (dpf). Additionally, we used the citrullination assay on lysates from whole 2 dpf larvae to detect calcium-dependent citrullination activity. Importantly, *padi2*^−/−^ zebrafish lysates lacked citrullination activity, even in the presence of excess calcium (Fig 1 G), indicating that Padi2 is likely the only protein with detectable citrullination activity in long pec zebrafish larvae. Interestingly, although previous studies have indicated that mammalian PAD1 and PAD6 are necessary for normal development and fertility (Esposito et al., 2007; Kan et al., 2012; Zhang et al., 2016), we found that the Padi2-deficient zebrafish did not display any gross morphological defects, had normal viability and crossings of this mutant line following expected Mendelian ratios (Fig S2 A and B). A homozygous incross produces viable and developmentally normal maternal-zygotic embryos indicating a maternal *padi2* contribution is not necessary during early embryonic development. To further address the role of citrullination in early development, we detected citrullination activity in cleavage- and gastrula-stage embryos and showed citrullination activity and Padi2 protein expression during both pre- and post- maternal to zygotic transition in wild-type embryos activity in 1-2 hpf larvae. This detected activity and protein was absent in the Padi2-deficient larvae (Fig 1 H and I). These observations provide the first evidence that citrullination is not necessary for broadly normal development in zebrafish.

The mammalian PAD2 is the predominant isozyme in skeletal muscle and nervous system (Kubilus and Baden, 1983; Watanabe et al., 1988; Watanabe and Senshu, 1989). To further characterize the mutant line, we examined the effects of the mutation on muscle development in zebrafish. We visualized slow and fast-twitch muscles in the trunk of 5 dpf larvae by staining for myosin heavy chain and F-actin. Both skeletal muscle fibers in the *padi2*^−/−^ larvae appeared morphologically comparable to wild-type (Fig S2 C and D). To examine neuromuscular synapses in the trunk of 5 dpf larvae, we immunostained for presynaptic vesicles (α-SV2) and acetylcholine receptors (AChR, α-BTX) (Fig S2 E and F). Quantification of these puncta showed that *padi2*^−/−^ larvae form more neuromuscular junctions than wild-type larvae (Fig S2 F). Previous studies show that Padi2 is expressed in central synapses (Bayes et al., 2017) and that PAD2 mice displayed behavioral defects (Falcao et al., 2019). These data suggest that citrullination may regulate the development of synapses, providing an interesting avenue for future investigation.

### Padi2 is required for efficient epimorphic regeneration

To determine the role of citrullination in wound healing and regeneration, we performed a tail transection of 2.5 dpf larvae through the notochord without wounding the caudal vein, as described by Rojas-Munoz *et al*. 2009. *padi2* expression during regeneration was determined at 24 hours post wounding (hpw) by qPCR analysis on extracted wounded fin tissue compared to age-matched, unwounded tissue (3 dpf). *padi2* was expressed in the affected tissue during regeneration and remained low in *padi2^−/−^* wounded fins (Fig S3 A). Regeneration of the fin was assessed 3 days post wounding (dpw) by measuring the fin length from the blood circulation loop to the edge of the fin along the notochord axis (Fig 2 A). Regrowth of the fin was impaired in the Padi2-decifient larvae compared to wild-type cousins (Fig 2 B). Similar effects were observed with transient morpholino depletion of *padi2* following a fin fold excision (Fig S3 B). *padi2^−/−^* larvae had a slight, but statistically significant, increase in their developmental fin length at 5 dpf compared to wild-type cousins (Fig 2 C), indicating Padi2 has different roles during fin development and regeneration. These findings suggest that Padi2 is necessary for efficient tail fin regeneration, identifying a new role for citrullination in wound repair. Interestingly, PAD activity and excessive citrullination has previously been linked with poor wound healing in mice and chick embryos (Coudane et al., 2011; Lange et al., 2011; Wong et al., 2015). It is possible that the absence of a PAD4 orthologue in zebrafish may contribute to these differences and their high regenerative capacity. Taken together, these findings support the idea that tight regulation of citrullination activity is necessary for normal regeneration. Furthermore, these results highlight the different mechanisms underlying developmental and regenerative growth.

**Figure 2:**
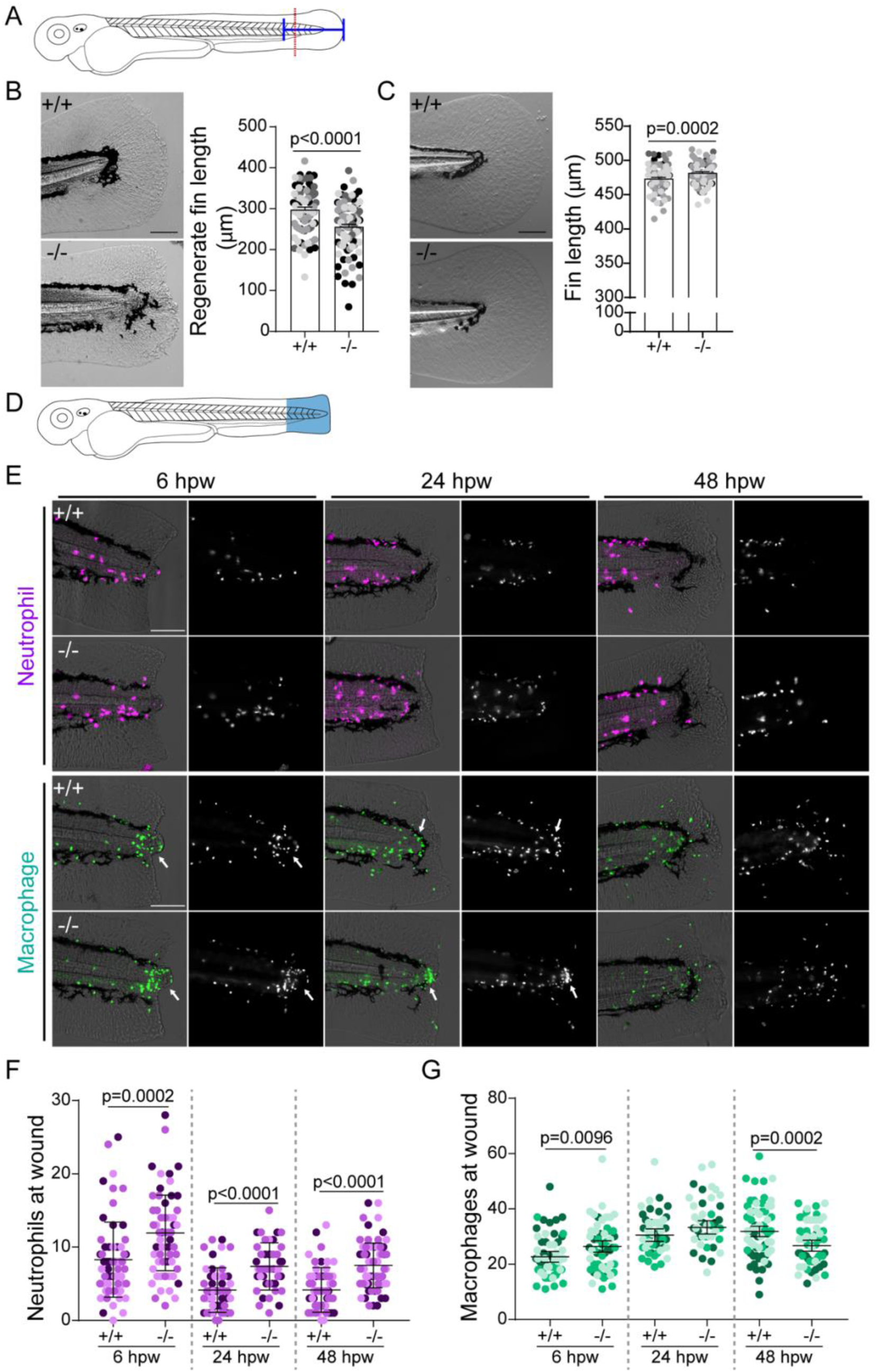
Padi2 is required for proper regeneration and leukocyte recruitment. (A) Schematic of regeneration assay. Tail transections were performed through the notochord (red dotted line) at 2.5 dpf. Fin lengths were measured from the blood circulation to the end of the fin (blue solid line). (B) Representative bright field images of regeneration at 3 dpw and quantification of regenerate fin length from 4 independent replicates with n = 90 +/+, 95 -/-. (C) Representative images of 5 dpf developmental fins and quantification of developmental fin length from 5 independent replicates with n = 109 +/+, 108 -/-. (D) Schematic of leukocyte quantification region (in blue) at a wound. (E) Representative images of leukocytes at a wound at 6, 24, and 48 hpw visualized with mCherry-labeled neutrophil nuclei (*Tg(lyzC:H2B-mCherry)*) and GFP-labeled macrophage nuclei (*Tg(mpeg1:H2B-GFP)*). Fluorescence channel on the right, merge with bright-field on the left. Macrophage localization to the periphery of the notochord bead indicated with an arrow. (F) Quantification of number of neutrophil nuclei at a wound from 3 independent replicates (6 hpw, n = 62 +/+, 57 -/-; 24 hpw, n = 50 +/+, 47 -/-; 48 hpw, n = 63 +/+, n=65 -/-). (G) Quantification of number of macrophage nuclei at a wound from 3 independent replicates (6 hpw, n = 61 +/+, 55 -/-; 24 hpw, n = 48 +/+, 44 -/-; 48 hpw n = 63 +/+, 57 -/-). All quantifications have lsmeans (± SEM) reported with p values calculated by ANOVA. Scale bars = 100 μm.

### Padi2 modulates leukocyte recruitment to a wound

A hallmark of wound repair is leukocyte infiltration and subsequent resolution of inflammation that can modulate wound healing (Wilgus et al., 2013). Citrullination has been shown to affect the immune response in human disease (Li et al., 2010), with direct evidence for deimination of leukocyte chemotactic cues (Loos et al., 2009; Proost et al., 2008; Yoshida et al., 2014). To visualize leukocyte responses to a wound, we compared *padi2^−/−^* and wild-type cousin larvae with either labeled neutrophils (*Tg(lyzc:H2B-mCherry)*) or macrophages (*Tg(mpeg1:H2B-GFP)*) and quantified leukocyte numbers within the region posterior to the blood circulation loop (Fig 2 D-G). Padi2-decifient larvae had a consistent increase in neutrophils at the wound at 6, 24, and 48-hour post wounding (hpw) (Fig 2 E and F). This difference is not due to altered total neutrophil numbers as total numbers in *padi2* mutants were not significantly different than their wild-type cousins (Fig S3 C), although there was a small increase of neutrophils in the unwounded fin (Fig S3 D). We also found a small increase in macrophages at the wound site in *padi2^−/−^* larvae, although this difference did not persist and there was no change in total macrophage numbers (Fig 2 E, G and Fig S3 E, F). Interestingly, macrophages displayed a localized aggregation around the notochord bead at 6 hpw and 24hpw (Fig 2 E), suggesting a potential role for macrophages in the notochord bead during wound healing. Taken together, these findings are consistent with a recent report showing that lymph nodes from PAD2 knockout mice show an increase in the expression of genes involved with leukocyte migration (Liu et al., 2018). It is unclear if the persistent leukocyte infiltration is due to a failure of wound resolution or due to a direct effect of citrullination on leukocyte signaling pathways. Citrullination of chemokines have been reported to dampen inflammatory signaling (Loos et al., 2008; Proost et al., 2008; Struyf et al., 2009), and the neutrophil chemokine, Cxcl8, and its receptors, Cxcr1 and Cxcr2, regulate neutrophil forward and reverse migration in response to a wound (Powell et al., 2017). Alternatively, citrullination of extracellular matrix (ECM) components, such as collagen or fibronectin, affect cell migration (Shelef et al., 2012; Sipila et al., 2014; Yuzhalin et al., 2018), and could potentially regulate inflammation by altering the wound ECM.

### Wounding induces localized histone citrullination in the notochord bead

We next considered whether wounding induces localized citrullination of histones in larval zebrafish due to the reported role of citrullinated histones in maintaining pluripotency (Christophorou et al., 2014; Wiese et al., 2019; Xiao et al., 2017). First, we assayed for total histone H4 citrullination, using immunoblotting, in whole larvae lysates treated *ex vivo* with calcium. Whole larvae lysate from wild-type larvae showed calcium-dependent citrullination of histone H4 (H4cit3) that was not present in *padi2^−/−^* lysate (Fig 3 A), indicating that Padi2 mediates histone citrullination in zebrafish larvae. While PAD4, but not PAD2 is required for histone citrullination in activated neutrophils in mammals (Holmes et al., 2019), our data suggest that zebrafish Padi2 adopts some functions of the later evolved PADs. Caudal fin transection results in increased intracellular calcium at a wound (Yoo et al., 2012a) which may promote citrullination. Visualization of histone H4 citrullination upon caudal fin amputation in wild-type zebrafish revealed signal exclusively within a localized group of cells in the notochord bead (Fig 3 B and C), a region previously described as the regeneration blastema (Rojas-Munoz et al., 2009). Immunofluorescence microscopy revealed H4cit3 signal as early as 1 hpw and this citrullination persisted up to 24 hpw (Fig 3 C). Histone H4 citrullination was diminished by 48 hpw, and was undetectable in the regenerated fin at 72 hpw (Fig 3 B and C). This histone H4 deimination is wound dependent as no signal was observed in unwounded larvae (Fig S3 G).

**Figure 3:**
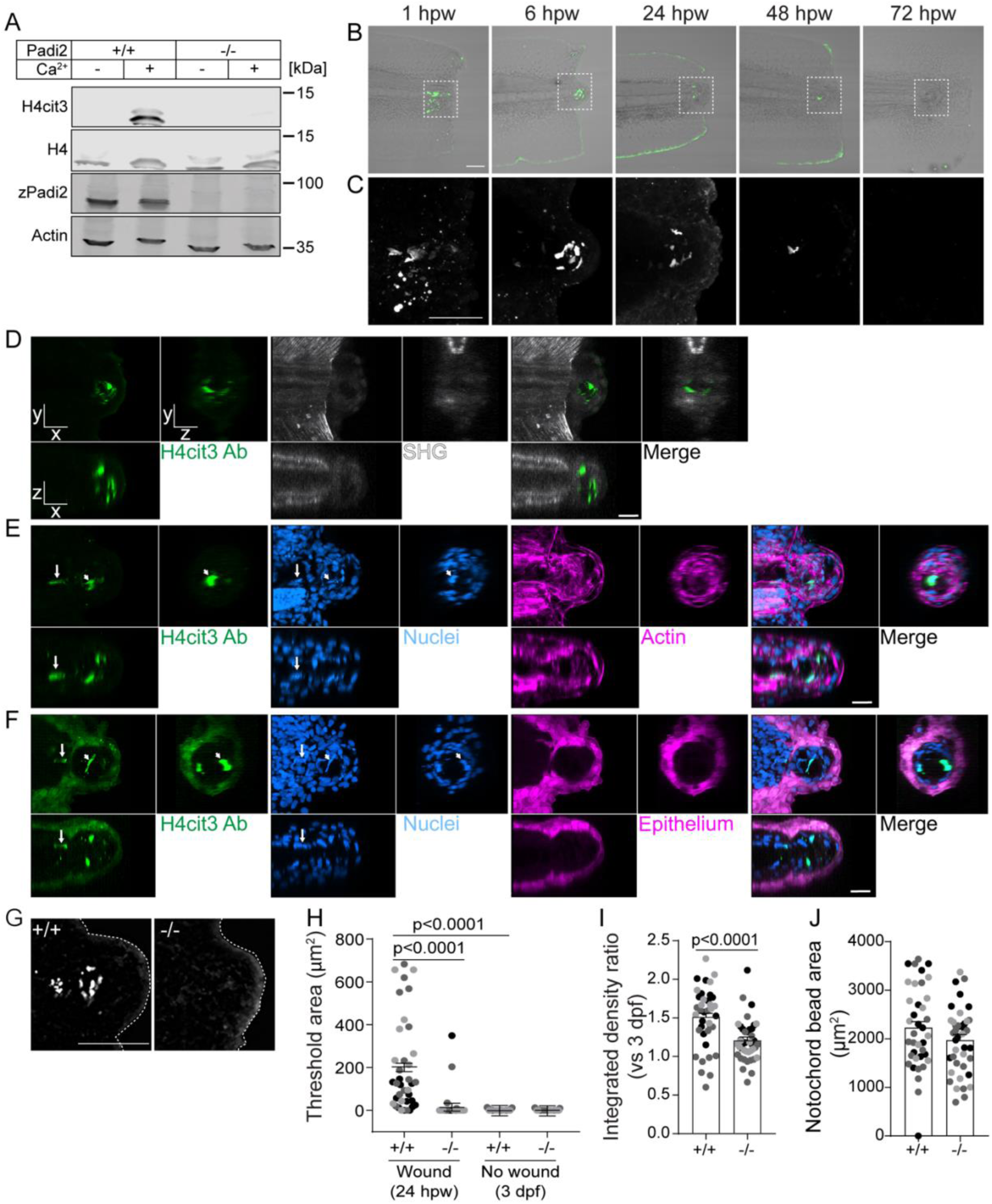
Tail transection stimulates localized Padi2-dependent histone H4 citrullination. (A) Representative western blot of *padi2^−/−^* and wild-type whole larvae lysates showing protein levels of: citrullinated Histone4 (H4cit3), total Histone4 (H4), total Padi2 (zPadi2), and total Actin (Actin) (representative of 2 replicates). (B) Representative images of H4cit3 antibody staining in tail transected fins at 20x magnification. Merge images with bright-field are shown for orientation. Box denotes region imaged at 60x magnification in (C) showing H4cit3 immunolabel alone. B-C, Representative images from 3 independent replicates. (D-F) Representative multiphoton microscopy enface (x,y view) and orthogonal (x, z view is below; y, z view is to the right) sections of 6 hpw wild-type caudal fins labeled with H4cit3 immunofluorescence (green) in conjunction with either (D) SHG (white) or (E) DAPI labeled nuclei (blue) and Rhodamine-phallodin labeled actin (magenta) or (F) DAPI labeled nuclei (blue) and (T*g(krt-4:EGFP)*) expressing epithelium (magenta). Note in (F) fluorophore tag for epithelium crosses into H4cit3 antibody channel at the setting needed to detect the antibody signal. Arrow indicates nucleus with citrullinated histones in the notochord region while arrowhead points to one example of a nucleus with citrullinated histones in the notochord bead. For image presentation, section thickness shown 2 μm for both x and y, 10 μm in z. Representative images from 2 independent replicates. (G) Representative images of H4cit3 immunostaining in 24 hpw wild-type cousin (left) and *padi2*^−/−^ (right). (H) Quantification of H4cit3 signal area at the notochord at 24 hpw and 3 dpf (no wound control). (I) Quantification of H4cit3 integrated density in 24 hpw larvae normalized to the average of 3 dpf for each genotype. Data are from 3 pooled independent (J) Quantification of the notochord bead area at 24 hpw from 3 pooled independent replicates. All quantifications have the lsmeans (±) SEM reported and p values calculated by ANOVA (H-J: 24 hpw, n = 38 +/+, 41 -/-; 3 dpf n = 25 +/+, 25 -/-). Scale bars = 100 μm in B, G; 50 μm in C; 20 μm in D, E, F.

To further characterize this structure, we used multiphoton microscopy to understand the 3D localization of citrullinated histones within the context of the wounded fin. Using second harmonic generation (SHG) to visualize the collagen fiber network, in conjunction with H4cit3 immunostaining, we observed citrullinated histones in a region devoid of collagen fibers at 6 hpw (Fig 3 D). The notochord bead containing this signal formed posterior to the original wound axis demarcated by the end of the collagen network. Labeling F-actin to visualize cell borders demonstrated that the notochord bead is composed of multiple cells (Fig 3 E). The H4cit3 signal colocalized with DAPI but only a subset of nuclei in this region were positive for histone citrullination. Nuclei with histone citrullination can also be observed in cells outside the notochord bead; it is possible that these cells move into this region from the notochord (Fig 3 E, F). While mammalian PAD2 lacks a canonical nuclear localization signal, evidence for nuclear localization and activity have been reported (Cherrington et al., 2010; Cherrington et al., 2012; Zheng et al., 2019). Future experiments will be needed to verify the mechanism by which zebrafish Padi2 translocates into the nucleus. Finally, visualization of the epithelium using Tg*(krt4:EGFP)* revealed that histone citrullination did not occur within epithelial cells (Fig 3 F). By 6 hpw, the wound epithelium has already formed and encompasses the citrullinated cells and notochord bead (Fig 3 F). Taken together, we have identified a new wound-induced structure within the notochord bead comprised of a subpopulation of cells with citrullinated histones. Given the previous work implicating citrullinated histones in pluripotency, this raises the intriguing possibility that these cells represent a key signaling hub in regenerative growth. Future work will be needed to identify which cells or signals are necessary to promote citrullination only within this subpopulation of notochord bead cells.

Toward this goal, we found that wound-induced histone citrullination was absent in *padi2^−/−^* larvae at 24 hpw (Fig 3 G). In the *padi2^−/−^* larvae, wounding did not induce histone H4 citrullination in the notochord bead above unwounded levels, in contrast to their wild-type cousins (Fig 3 G-I). Morphologically, both wild-type and Padi2-deficient larvae formed similar sized notochord beads early after wounding (Fig 3 J). Previous reports show that this region of cells act as a required wound-signaling center and blastema structure that orchestrates regeneration (Rojas-Munoz et al., 2009; Romero et al., 2018). Importantly, this population of cells with citrullinated histones were associated with the blastema reporter, *Tg(lepb:EGFP)* (Kang et al., 2016) (Fig S3 H and I). Our data indicate that citrullination is necessary for efficient regeneration and that a localized population of cells within the blastema structure contains citrullinated histones. With a known role for histone citrullination in stem cell maintenance, it is intriguing to speculate that this population of cells with citrullinated histones has pluripotent features required for efficient tissue repair by acting as a multipotent signaling center.

### Padi2-deficient larvae have impaired wound-induced proliferation

An essential aspect of epimorphic regeneration is remodeling by wound-induced apoptosis and proliferation (Gauron et al., 2013; Nechiporuk and Keating, 2002; Tseng et al., 2007). We did not observe a significant change in wound-stimulated apoptosis in *padi2* mutant larvae (Fig S3 J-L). We assayed cell proliferation using EdU incorporation and found that mutant larvae had a greater number of EdU-positive cells within the developing caudal fin than wild-type larvae (Fig 4 A and B), consistent with the observed increase in *padi2^−/−^* developmental fin size (Fig 3 C). Upon wounding, wild-type larvae had an almost 4-fold increase in proliferative cells within the regenerating fin compared to the unwounded fins. By contrast, Padi2-decifient larvae exhibited impaired induction of proliferation, with only a 2-fold induction of proliferation after wounding (Fig 4 C-E). Similarly, *padi2* morpholino knockdown resulted in decreased mitotic index at 24 hpw compared to control injected embryos (Fig S3 M and N). To further quantify wound-induced proliferation, we focused on the dorsal region of the tail since much of the developmental proliferation is localized to the ventral fin. In this dorsal region we observed impaired proliferation in the *padi2^−/−^* larvae compared to wild-type cousins (Fig 4 F and G). These findings suggest opposing roles for Padi2 in developmental fin and wound-induced proliferation, supporting the idea that these two processes have distinct modes of regulation.

**Figure 4:**
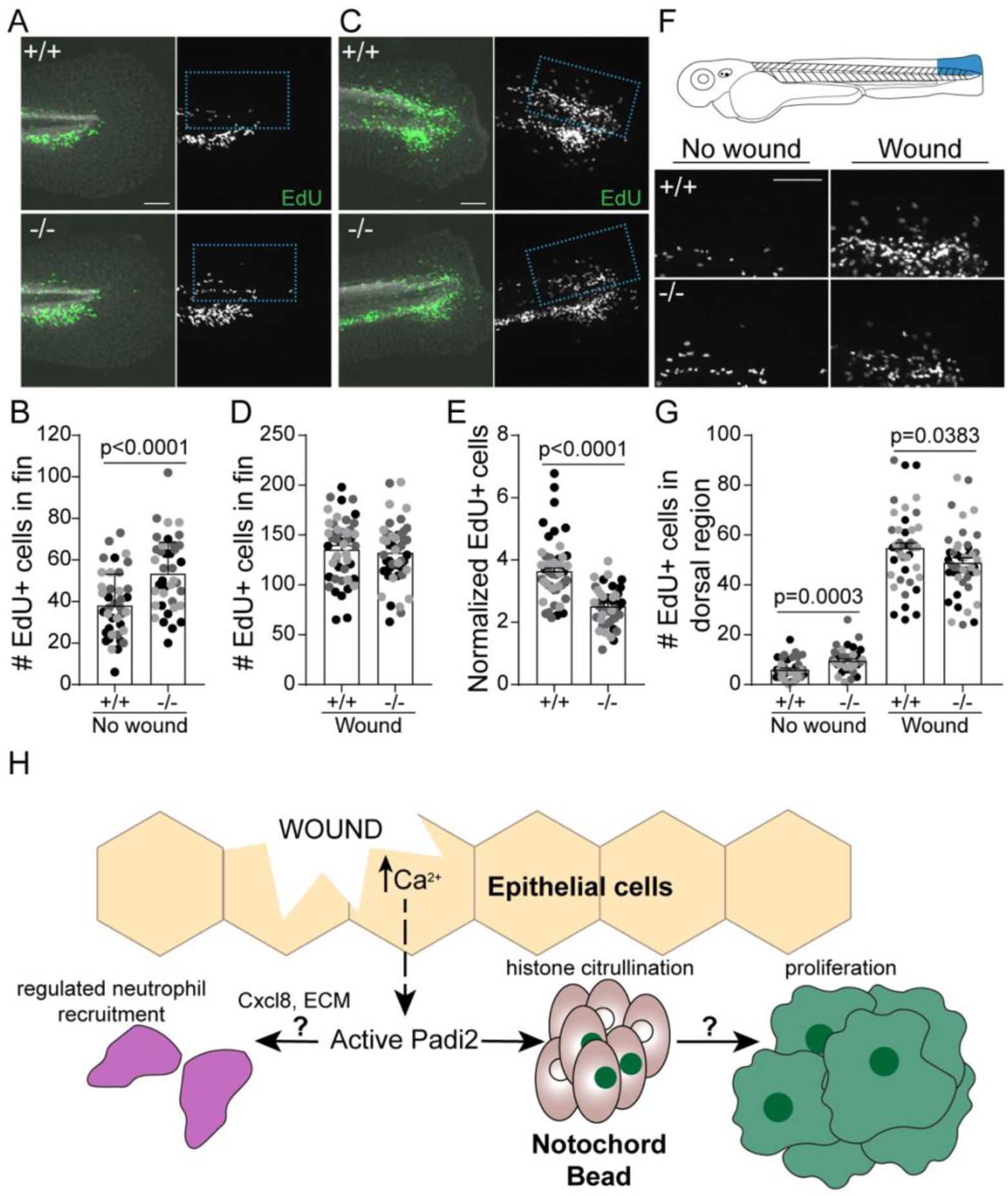
Wound-induced proliferation is perturbed in Padi2-deficient larvae. (A,C) Representative images of 6-hour EdU pulsed larvae in (A) developmental, unwounded or (C) 66 hpw fins. Merged images of EdU (green) and DAPI (white) on the left and single EdU (white) image on the right. (B,D) Quantification of EdU-positive cells in the fin. (E) Number of EdU-positive cells in the fin normalized to corresponding no wound conditions. (F) Representative images of the dorsal half of 6-hour EdU-pulsed fins. (G) Quantification of EdU-positive cells within the dorsal region of the fin. All data are from 3 pooled independent replicates with the lsmeans and SEM reported and p values calculated by ANOVA (no wound, n = 39 +/+, 39 -/-; 66 hpw n = 47 +/+, 47 -/-). Scale bars = 100 μm. (H) A proposed model depicting how the early wound epithelial calcium flux might activate (dashed arrow) Padi2 to catalyze citrullination events that, either directly or indirectly (left question mark), regulate neutrophil (purple) recruitment to the wound, possibly mediated by the Cxcl8 pathway or ECM modification. Concomitantly, wound-dependent Padi2 citrullination of histones (green nuclei) within a subset of cells in the notochord bead (pink) potentially stimulates, through yet to be determined mechanisms (right question mark), regenerative proliferation.

In summary, we identified a new role for citrullination in wound healing and regeneration. Early calcium flux is a universal injury signal in organisms ranging in complexity, and while there are many citrullination-independent calcium-induced wound pathways (Niethammer, 2016), this work identifies one potential regenerative mechanism downstream of early wound-induced calcium flux (Fig 4 H). We showed that zebrafish Padi2 has conserved activity and calcium dependence, and that it is necessary for calcium-mediated histone citrullination in zebrafish larval lysates. Padi2 appears to have opposing roles in developmental fin growth and regeneration with respect to proliferation and tissue growth and is required for proper neutrophil response to a wound. The identification of a new, localized population of cells with specifically wound-induced, Pad2-dependent histone citrullination in the notochord bead suggests that citrullination in this region may play a key role in orchestrating efficient regenerative growth. The known role of histone citrullination in gene expression and pluripotency suggests that this small population of cells represents a unique subset of blastemal cells. A future challenge will be to characterize these cells further including analysis of their gene expression profile to identify specific downstream effectors of this localized, regulated histone citrullination. Moreover, this citrullination-deficient vertebrate model provides a powerful tool for future studies to dissect the role of citrullination in development, disease, and wound healing, and will aid in the identification of *in vivo* Padi targets.

## Materials and methods

### Zebrafish maintenance and handling

All protocols in this study were approved by the University of Wisconsin-Madison Animal Care and Use Committee (IACUC). Adult zebrafish were maintained on a 14h:10h light/dark schedule. Fertilized embryos were transferred and maintained in E3 buffer at 28.5°C. This study utilized adult AB and NHGRI-1 (LaFave et al., 2014) fish (obtained from the Zebrafish International Resource Center (ZIRC)) as well as previously published transgenic lines *Tg(mpeg1:H2B-GFP)* (Miskolci et al., 2019), *Tg(lyzc:H2B-mCherry)* (Yoo et al., 2012b), *Tg(krt4:EGFP)* (Yoo et al., 2012a), and *Tg(lepb:EGFP)* (Kang et al., 2016).

### Zebrafish and Human PADI alignment

Sequence alignments were performed using the EMBOSS Water pairwise sequence alignment algorithm (Smith and Waterman, 1981). Predicted transcripts are listed in Table 1. Transcript annotations are from GRCz10 with transcript 201a indicating a sequence slightly divergent from GRCz10’s transcript 201.

**Table 1.** Annotated PADI transcripts used in this study.

| Name | Transcript ID | Genome assembly |
| --- | --- | --- |
| zPadi2 202 | ENSDART00000140943.3 | GRCz11 |
| zPadi2 203 | ENSDART00000140943.2 | GRCZ10 |
| zPadi2 201 | ENSDART00000064842.6 | GRCz11 |
| zPadi2 201 | ENSDART00000064842.5 | GRCz10 |
| zPadi2 202 | ENSDART00000127766.2 | GRCz10 |
| hPADI2 202 | ENST00000375486.8 | GRCh38 |

### Generation of a padi2 mutant line and genotyping

Zebrafish CRISPR/Cas9 injections were performed as previously described in our lab (LeBert et al., 2018; LeBert et al., 2015). Guide RNA (gRNA) for zebrafish Padi2 (ENSDARG00000044167) was designed using CHOPCHOP (Montague et al., 2014). Exon 7 target sequence: GGGAACAGACACGCTGACGC

The pT7 gRNA was prepared as previously described (LeBert et al., 2018). The gRNA and Cas9 protein (New England Biolabs) were injected into one-cell stage NHGRI-1 embryos in a 2 nl volume consisting of ∼50 ng/μl gRNA and ∼300 nM Cas9. To confirm genome editing by the gRNA, genomic DNA was extracted from 2 days post fertilization (dpf) embryos, amplified using the primers listed below, and separated on a 3% MetaPhor gel (Lonza).

Padi2 F: CTGATACATGGCACAACCTACG

Padi2 R: GAAAACCAGCAAGCAGAGAAAGTT

Sequences of F0 mosaic cuts were confirmed by TOPO cloning (Invitrogen) and sequencing. Clutches of larvae with confirmed CRISPR cuts were grown to adulthood. Adult F0 CRISPR injected fish were screened for germline mutations by testing their individual outcrossed offspring (2-5 dpf) using the primers listed above and Indel Detection and Amplicon Analysis (IDAA) (Yang et al., 2015). Sequences were analyzed using Peak Studio (McCafferty et al., 2012). Mutation sequence was confirmed by TOPO cloning and sequencing.

Heterozygous *padi2* zebrafish were maintained by outcrossing the CRISPR mutants to AB wild-type background zebrafish and genotyped by genomic DNA isolated from fin clips and amplified using the primers listed above. PCR product was either separated on a 2% agarose gel for 3 hours or digested overnight with MwoI (New England Biolabs) and separated on an agarose gel to determine individual fish genotypes. For experimental purposes, F2 or F3 heterozygotes were incrossed for the generation of adult homozygous mutant and wildtype siblings. These adults were then incrossed to produce *padi2^−/−^* and wild-type clutches, referred to as cousins, which were only used for experimentation and not for the maintenance of subsequent generations.

### qRT-PCR

RNA and DNA were extracted from individual 2 dpf embryos from a *padi2*+/− incross using TRIZOL (Invitrogen) following the manufacturer’s protocol. Embryos were genotyped using GoTaq (Promega) as described above and 2-3 embryos of each genotype were used for cDNA production using Superscript III First Strand Synthesis System with Oligo(dT) (Thermo Fisher Scientific). qPCR was performed using FastStart Essential Green Master (Roche) and a LightCycler96 (Rocher). Primers for *padi2* and *ef1a* are listed below. Data were normalized to *ef1a* using the ΔΔCt method (Livak and Schmittgen, 2001) and represented as fold change over wild-type embryos.

For evaluation of *padi2* mRNA expression during wounding, incrosses of F3 or F4 adult wild-type and *padi2^−/−^* siblings were done to produce offspring cousins homozygous for the *padi2* mutation or wild-type. Fin samples were amputated at the line of the blood circulatory loop and 50-100 fins were pooled and flash frozen, with equivalent sample sizes used per replicate. RNA was extracted from fin tissue from 24 hpw and unwounded, 3 dpf, larvae, as described above. Primers for *padi2* and *rps11* (de Oliveira et al., 2013) are listed below. Data were normalized to *rps11* and represented as fold change over wild-type, unwounded, age-matched control fins.

Padi2 exon5 qRT-PCR F: TAATGGCCATGGTGCAGTTC

Padi2 exon6 qRT-PCR R: ATGGTCCATTAGTGCGCAAC

Ef1a qRT-PCR F: TGCCTTCGTCCCAATTTCAG

EF1a qRT-PCR R: TACCCTCCTTGCGCTCAATC

Rps11 qRT-PCR F: TAAGAAATGCCCCTTCACTG

Rps11 qRT-PCR R: GTCTCTTCTCAAAACGGTTG

### Generation of zebrafish *padi2* clones and point mutations

Padi2 splice variants were amplified with Pfu Turbo DNA polymerase (Agilent) from cDNA using In-Fusion primers listed below. PCR products and a pCS2+8 vector (Gokirmak et al., 2012) (Addgene) were digested with XbaI and BamHI (Promega) and ligated at room temperature using Takara ligation kit for long fragments.

Padi2 cloning R, with XbaI: GGATCG TCTAGATTACAGCTCCAGGTTCCACC

Padi2 cloning F transcript 201: CGATCCGGATCCATGGTGTCCCGTCGATCTCTTAC

Padi2 cloning F transcript 202: CGATCCGGATCCATGAATGTTTCGCAGGAGC

Both cDNA transcripts were cloned into pTRCHisA vector (Invitrogen) for N-terminal polyhistidine (his) tagging and expression in *E. coli* (BL21(DE3)pLysS Competent cells) using primers listed below. Constructs were inserted into the vector cut with BamHI and HindIII (Promega) using In-Fusion HD cloning kit (Clonetech). Point mutations were made with complementary primers (listed below) in pTRCHisA-padi2 vectors using QuikChange II Site-Directed Mutagenesis Kit (Agilent).

Catalytic C→A F: GTGAAGTTCACGCCGGGTCCAATGTTC

Catalytic C→A R: GAACATTGGACCCGGCGTGAACTTCAC

Ca1 binding Q→A F: ATCGCTGGATGGCGGATGAGCTTGAGTT

Ca1 binding Q→A R: AACTCAAGCTCATCCGCCATCCAGCGAT

Ca1 binding E→A F: GGATGAGCTTGCGTTTGGTTACATTG

Ca1 binding E→A R: CAATGTAACCAAACGCAAGCTCATCC

Ca1 binding E→A F: TTTCGGTAATCTGGCGGTCAGTCCACCA

Ca1 binding E→A R: TGGTGGACTGACCGCCAGATTACCGAAA

Ca2 binding D→A F: TGTTGTCCTGGCTTCTCCTCGTGAT

Ca2 binding D→A R: ATCACGAGGAGAAGCCAGGACAACA

### Antibody production and western blotting

The anti-zebrafish Padi2 antibody was generated in rabbits using combined full length 201a and 202 variants fused to 6x poly-histidine in the pTRCHisA vector. Each immunogen was purified from BL21 *E. coli* lysates using a nickel-nitrilotriacetic acid superflow resin (Qiagen) then combined and sent for anti-sera production (Covance). For western blotting, 50-100 ∼2 dpf or 5 dpf larvae were pooled and deyolked in calcium-free Ringer’s solution via gentle disruption with a p200 pipette. Lysates from 2 hpf and 7 hpf larvae were not deyolked; samples were instead dechorionated on a petri dish coated with 2% agarose and then rinsed with PBS. Larvae were washed twice with phosphate-buffered saline (PBS) and stored at −80°C until samples were lysed by sonication in 20mM Tris pH 7.6, 0.1% Triton-X-100, 0.2 mM phenylmethylsulfonyl fluoride (PMSF), 1 μg/mL Pepstatin, 2 μg/mL Aprotinin, and 1 μg/mL Leupeptin at 3 μL per larvae while on ice and clarified by centrifugation. Protein concentrations were determined using a bicinchoninic acid protein assay kit (Thermo Fisher Scientific), according to the manufacturer’s instructions. Equal amounts of total protein were loaded on 6-20% gradient SDS-polyacrylamide gels and transferred to nitrocellulose. For citrullination analysis by western of whole zebrafish lysates, methods for the citrullination colorimetric assay were followed, as described below, with the addition of dilution buffer in place of BAEE (N_α_-Benzoyl-L-arginine ethyl ester hydrochloride in 100 mM Tris pH 7.4). The reaction was stopped after 90 minutes by boiling samples in SDS-PAGE sample buffer. zPadi2 rabbit anti-serum was used at 1:500 dilution, anti-Histone H4 (citrulline 3) (EMD-Millipore) at 1:50, anti-actin (ac15; Sigma) at 1:1000, and anti-Histone H4 (EMD-Millipore) at 1:1000. Western blots were imaged with an Odyssey Infrared Imaging System (LI-COR Biosciences).

### *padi2* mRNA re-expression

*padi2*-201a cloned into pCS2+8 (described above) was linearized using NotI restriction digest and RNA was *in vitro* transcribed using the mMessage mMachine Sp6 kit (Ambion). RNA was cleaned up using an RNeasy Minikit column (Qiagen) and injected into single cell embryos (3nl of 100ng/ul). Embryo lysates were collected as described for western blotting at 2dpf and 5dpf.

### *In vitro* citrullination colorimetric assay

Zebrafish Padi2 constructs and point mutations were expressed in BL21 *E. coli* cells. Lysates were prepared on ice by sonication in 20mM Tris pH 7.6, 0.1% Triton-X-100, 0.2 mM phenylmethylsulfonyl fluoride (PMSF), 1 μg/mL Pepstatin, 2 μg/mL Aprotinin, and 1 μg/mL Leupeptin and clarified by centrifugation. Bacterial lysates were aliquoted and frozen at −80°C. Lysates from zebrafish larvae were prepared as described above for western blotting and used at equivalent amounts. Assay performed as previously described (Nakayama-Hamada et al., 2005). In short, 12.5μL lysate was incubated with 12.5 μL 4X reaction buffer (400 mM Tris pH 7.4, ± 80 mM CaCl_2_, 20 mM DTT), 12.5 μL 80 mM BAEE (N_α_-Benzoyl-L-arginine ethyl ester hydrochloride in 100 mM Tris pH 7.4), 12.5 μL dilution buffer (10 mM Tris pH 7.6, 150 mM NaCl, 2mM DTT) for 1 hour at 37°C. Reaction was stopped by the addition of 33 μM EDTA final concentration. Reactions were diluted 1:10 for an 8 mM BAEE final concentration and 50 μL aliquots were done in triplicate in a 96-well plate. 150 μL colorimetric buffer (composed of 1 mL buffer A (80 mM diacetyl monoxime, 2 mM thiosemicarbazide) and 3 mL buffer B (3 M phosphoric acid, 6 M sulfuric acid, 2 mM ammonium iron (III) sulfate)) were added to each well and incubated at 95°C for 15 minutes, absorption was read at 540 nM. Relative light units were normalized to western blot densitometry using Odyssey Infrared Imaging System (LI-COR Biosciences).

### Morpholino injections

Morpholino oligonucleotides (Genetools) were designed to the intron1/exon2 border of *padi2*. Morpholinos were resuspended in water to a final concentration of 1 mM. Morpholinos were diluted to a final concentration of 100 μM and 3 nl injection mix was injected into one-cell stage embryos which were subsequently maintained at 28.5°C. Morpholino sequences used: *padi2* MO: 5’-GAGCACATCTGGAATGGGAATATAT; control MO: 5′-CCTCTTACCTCAGTTACAATTTATA-3′.

### Regeneration assays

For larval regeneration assays, incrosses of F3 or F4 adult wild-type and *padi2^−/−^* siblings were done to produce offspring cousins homozygous for the *padi2* mutation or wild-type. Dechorionated larvae were transferred to 35 mm milk-coated plates. Larvae were washed twice in E3 and wounded in a final 0.24 mg/mL tricaine (ethyl 3-aminobenzoate, Sigma)/E3 solution. Tail transections were performed on ∼2.5 dpf larvae with a surgical blade (feather no 10) roughly four vacuolated cells from the posterior end of the notochord. Larvae were again washed 3 times with E3 and allowed to regenerate for 3 days post-wounding (dpw), at which point larvae were fixed with 4% paraformaldehyde (PFA; Sigma-Aldrich) in PBS at 4°C overnight. Fins were imaged on a Zeiss Zoomscope (EMS3/SyCoP3; Zeiss; 1x Plan-NeoFluor Z objective) with an Axiocam Mrm CCD camera using ZEN pro 2012 software (Zeiss). Regenerate length was measured from the edge of the blood vessel to the caudal edge of the tail fin using the FIJI image analysis software (Schindelin et al., 2012)). Unwounded, 5 dpf larvae fin lengths were measured as a developmental control. Fin transections were performed on MO injected larvae similarly to as described above with amputation adjacent to the notochord, without causing damage to the notochord. Regenerated fins and developmental controls were measured from the caudal tip of the notochord to the caudal edge of the tail fin.

### Immunofluorescence, microscopy and analysis

Images always shown with anterior to the left.

#### Neuromuscular labels

Immunostaining was performed on cousin offspring from incrossed adult F2 wild-type siblings and incrossed *padi2^−/−^* sibling zebrafish. 5 dpf larvae were fixed in 4% PFA, 0.125 M sucrose, and 1X PBS overnight at 4°C. For detection of slow muscles, larvae were washed 3 times with 0.1% PBS-Tween20 and incubated in 0.1% w/v collagenase type 1A (Sigma) in PBS at 37°C for 1.5 hours, followed by 3 washes in PBSTD (0.3% TritonX, 1% DMSO in PBS). Larvae were blocked for 2 hours at room temperature (RT) in PBSTD with 2% BSA and 4% goat serum. Monoclonal mouse anti-myosin heavy chain antibody (F59) (DSHB) (Miller et al., 1985) was used at 1:20 in block buffer and incubated overnight in 4°C. Larvae were washed five times in PBSTD and secondary Dylight 488 donkey anti-mouse IgG antibody (Rockland Immunochemicals) was used at 1:250 in block buffer overnight at 4°C. Final five washes were done in PBSTD. Images were acquired on a spinning disk confocal (CSU-X; Yokogawa) on a Zeiss Observer Z.1 inverted microscope and an EMCCD evolve 512 camera (Photometrics) with a Plan-Apochromat NA 0.8/20x air objective and collected as z-stack of 1 μm optical sections at 512×512 resolution. Images were z-projected with using Zen 2.3 lite software (Zeiss).

For visualization of fast muscle, fixed fish were washed with PBS 3 times followed by three washes in PBS with 0.1% Tween20. Larvae were permeabilized with PBS 2% PBSTx (20% Triton-X-100 in 1X PBS) for 1.5 hours with gentle rocking. Fish were then incubated with Rhodamin-phalloidin (Invitrogen) diluted 1:100 in 2% PBSTx at 4°C overnight. Fish were rinsed in fresh 2% PBSTx followed by several washes in 0.2% PBSTx. Imaging was performed on the spinning disk microscope with a Plan-Apochromat NA 0.8/20x air objective (centered on cloaca) with 1 μm optical sections.

For neuromuscular junction visualization, fix was washed off with three PBS washes. The skin was peeled with fine forceps (Dumont #55 dumostar, Fine Science Tools) starting above the swim bladder and removed down to the fin. Skinned larvae were incubated in 0.1% w/v collagenase type 1A at RT for 15 minutes with gentle rocking followed by three washes in PBS. For detection of acetylcholine receptors (AChR), larvae were incubated for 30 minutes at RT 10 μg/ml Alexa 594 conjugated a-bungarotoxin (Thermo Fisher Scientific) diluted in incubation buffer (IB: 0.1% sodium azide, 2% BSA, 0.5% Triton-X-100 in PBS, pH7.4). Embryos were rinsed three times in IB.Mouse anti-synaptic vesicle glycoprotein 2A antibody (SV2) (DSHB) (Buckley and Kelly, 1985) was used at 1:50 in IB overnight at 4°C. Larvae were washed 5 times in IB and incubated with secondary Dylight 488 donkey anti-mouse IgG antibody (Rockland Immunochemicals) at 1:250 in IB for 4 hours at RT or 4°C overnight. Final washes were done in IB before imaging on a spinning disk microscope with an EC Plan-NeoFluaR NA 0.75/40x air objective (Zeiss) (centered around the cloaca with 2×1 tile images and 1 μm optical section z-stacks). To quantify colocalization of signal, maximum intensity projections were analyzed in FIJI using the plugin ComDet v3.7 for spot localization (https://github.com/ekatrukha/ComDet/wiki). Particles were threshold as approximate size being 5 pixels, intensity threshold for SV2 between 4-5 and α-BTX between 2-3 and a 6 pixel max distance between particles.

#### Histone citrullination

Immunostaining was performed on offspring cousins from incrossed adult F3 wild-type siblings and incrossed *padi2*^−/−^ siblings. To identify histone citrullination, larvae were fixed in a solution of 1% NP-40, 0.5% Triton-X, and 1.5% PFA in PBS at 4°C overnight. The following day fix was replaced with a block solution of 2.5% BSA, 0.5% Tween-20, 5% goat serum in PBS. Samples were blocked for at least 2.5 hours at room temperature followed by the addition of poly-clonal rabbit anti-histone H4 (citrulline 3) antibody (EMD Millipore) used at 1:100 and incubated overnight at 4°C. For time course experiments, samples were kept in block at 4°C until the final sample was prepared, at which time all samples were blocked at room temperature before the addition of the primary antibody. Samples were washed 3 times in PBS at room temperature for 5 minutes each and secondary Dylight 488 donkey anti-rabbit (Rockland Immunochemicals) or Alexa Fluor 568 goat anti-rabbit IgG antibodies (Invitrogen) were used at 1:250 in block buffer overnight at 4°C. When indicated, Rhodamine-phalloidin (Invitrogen) and 10 mg/mL DAPI (4’,6-diamidino-2-phenylindole; Sigma) were added with secondary antibodies at 1:100 and 1:10,000 dilutions, respectively. Finally, 4 washes were done in PBS. Images were acquired on a laser-scanning confocal microscope (FluoView FV1000; Olympus) with an NA 0.75/20x or PLANAPO NA 1.45/60x oil objective and FV10-ASW software (Olympus). 20x images used for quantification were acquired as Z-stacks with 25, 1 um optical slices at 640×640 resolution. Alternatively, images were acquired using multiphoton microscopy. For this, caudal fins of fixed, PTU-treated, labeled larvae were removed from the trunk with a scalpel blade (Feather #15) then imaged in a 50 mm coverglass (#1.5) bottom dish (MatTek, Ashland MA) in PBS, as previously described (LeBert et al., 2016; LeBert et al., 2015). A second coverslip over the glass bottom depression minimized sample movement. The fins were imaged on a custom-built multiphoton microscope (Conklin et al., 2011; LeBert et al., 2016) at the Laboratory for Optical and Computational Instrumentation using a 40X long working distance water immersion lens (1.2 NA, Nikon, Melville NY). All signals were detected sequentially using a H7422P-40 GaAsP Photomultiplier Tube (PMT) (Hamamatsu, Japan). The backwards SHG signal was collected with the multiphoton source laser (Chameleon UltraII, Coherent Inc., Santa Clara, CA) tuned to 890 nm, with a 445/20 nm bandpass emission filter (Semrock, Rochester NY). The fluorescent signal from H4cit3 antibody was collected using a either a 520/35 nm bandpass emission filter (Semrock) for the Dylight 488 donkey anti-rabbit secondary antibody (Rockland Immunochemicals) or a 615/20 nm bandpass emission filter (Semrock) for the Alexa Fluor 568 goat anti-rabbit secondary antibody (Invitrogen). The 615/20 emission filter was used to collect the fluorescent signal from the Rhodamine-Phalloidin while the 520/35 nm emission filter was used to detect the krt4:EGFP and lepb:EGFP fluorescence. DAPI fluorescence was excited with the laser tuned to 740 nm and the emission collected using the 445/20 filter. Brightfield images were simultaneously collected using a separate photodiode-based transmission detector (Bio-Rad, Hercules CA). Data were collected as z-stacks with optical sections 2 microns apart, at 512 x 512 resolution.

#### Mitotic index

For evaluation of cells undergoing mitosis, 24 hpw and 3 dpf MO-injected larvae were fixed with 1.5% PFA in 0.1M PIPES, 1.0 mM, 2 mM EGTA overnight at 4°C and immunolabeled with phosphorylated histone H3 (serine10) antibody (Millipore). To remove fixation solution, larvae were washed with PBS three times and placed in methanol at −20°C overnight. Samples were rehydrated in subsequent 5 minute washes at ratios of 2:1, 1:1, 1:2 methanol:PBSTx (PBS with 0.2% Triton-X), and a final PBSTx wash. Larvae were incubated in 0.15 M glycine in PBS for 10 minutes at room temperature followed by 3 PBSTx washes. Fish were blocked in 1% BSA in PBSTx for 1 hour at room temperature. Phosphorylated histone H3 (serine10) antibody diluted 1:300 in block was incubated overnight a 4°C. Samples were washed for 15-30 minutes in block, twice in PBSTx, and another wash in block. Incubation with Dylight donkey anti-rabbit 488 secondary was used, followed by four washes in PBSTx. Samples were imaged and quantified on the laser-scanning confocal microscope with a 20x lens, as described above.

### Leukocyte Imaging

*padi2^+/-^* adults were crossed to AB wild-type zebrafish labeled with macrophage nuclei (*Tg(mpeg1: H2B-GFP))* or neutrophil nuclei (*Tg(lyzc:H2B-mCherry*)) and subsequently incrossed to produce homozygous, fluorescently labeled adults. Experiments were performed on wild-type cousins and *padi2^−/−^* larvae resulting from incrossed adult transgenic siblings. Wounding was performed as described above and larvae were fixed with 1.5% PFA in 0.1 M PIPES (Sigma-Aldrich), 1 mM MgSO_4_ (Sigma-Aldrich), and 2 mM EGTA (Sigma-Aldrich) overnight at 4°C. Wounds were imaged on a Zeiss Zoomscope, as above. Leukocyte numbers were counted by hand in the region past the blood circulatory loop. Whole larvae images were acquired on a spinning disk confocal (CSU-X; Yokogawa) on a Zeiss Observer Z.1 inverted microscope and an EMCCD evolve 512 camera (Photometrics) with a Plan-Apochromat NA 0.8/20x air objective (5 μm optical sections, 5×1 tiles, 2355×512 resolution).

### EdU and apoptosis labeling

Immunostaining was performed on offspring “cousins” from incrossed adult F3 wild-type siblings and incrossed *padi2*^−/−^ siblings. Proliferation in the fin was measured using Click-iT Plus EdU Imaging Kit (Life Technologies). Larvae were incubated in 10 μM EdU (5-ethynyl-2’-deoxyurdine) solution in E3 for 6 hours with slight agitation. Wounded fish were incubated from 60-66hpw along with age matched unwounded controls. Larvae were fixed in 4% PFA in PBS overnight at 4°C and stored in methanol at −20°C until staining. Staining protocol was conducted according to manufacturer’s instructions. EdU-stained larvae were also incubated with rabbit anti-active Caspase3 antibody (BD Biosciences) at 1:200 in block (PBS, 1% DMSO, 1% BSA, 0.05% Triton-X, 1.5% goat serum) followed by incubation with Alexa 550 goat anti-rabbit secondary antibody and 0.01 mg/mL DAPI (Sigma). Immunofluorescence images were acquired on a spinning disk confocal (CSU-X; Yokogawa) on a Zeiss Observer Z.1 inverted microscope with an EMCCD evolve 512 camera (Photometrics) and a Plan-Apochromat NA 0.8/20x air objective, as Z-stacks, 3 μm optical sections, and with 512×512 resolution.

### Image analysis/processing

Image analysis was performed on FIJI. For experiments where fluorescence intensity was quantified, no adjustments were made to the images prior to analysis. For Histone H4cit3 analysis, a region of interest 92 x 93 microns was centered around the notochord, as determined by the corresponding bright-field image. Immunostained images were z-projected as a maximum intensity projection and the integrated density in the region of interest (ROI) was determined. Images were thresholded using the threshold plugin using auto-thresholding with the “Intermodes” method in Fiji (Prewitt and Mendelsohn, 1966) and the total area within the ROI was determined for particles larger than 8 pixels. For presentation purposes, images were processed to remove background using despeckling. Notochord bead area was determined in FIJI by outlining this structure as determined by examination of the optical bright-field slices.

Total neutrophil numbers were determined using Imaris (Bitplane) with the spots function as defined by a 10 μm diameter in the XY plane and a Z-diameter of 20 μm. Total macrophage numbers were counted by hand using Z-projected images in Zen 2.3 lite software. For total leukocyte quantifications, leukocytes within the yolk sac and heart were excluded.

For spatial assessment of nuclei with citrullinated histones, three dimensional reconstructions and slices were constructed using Imaris (Bitplane, Oxford Instruments, UK). Videos of z-stack scans and 3D rotations were made in Imaris, annotated in FIJI using “<u>Annotation_to_overlay1.3</u>” plugin (https://www2.le.ac.uk/colleges/medbiopsych/facilities-and-services/cbs/AIF/software-1/imagej-macros#Annotation) and converted to MP4 using HandBrake (v1.2.2) software (The HandBrake Team, https://handbrake.fr/).

For EdU analysis, images were 3D reconstructed using Imaris software (Bitplane). The number of EdU-positive cells were quantified in the fin region posterior of the blood circulatory loop with the spots function as defined by an XY-diameter of 7 μm and a Z-diameter of 14 μm. The level of apoptosis activation at the wound was determined by outlining the fin past the blood circulation using the corresponding bright-field image. In FIJI, total threshold area for active-Caspase3 signal in the wound was determined using the threshold plugin in Fiji by auto-thresholding with the “Yen Dark” method (Yen et al., 1995) for particles larger than 3 pixels.

### Statistical analysis

For all statistical analyses, at least three independent replicates were conducted. For data in Fig 1 G, analysis was done using one-way ordinary analysis of variance (ANOVA) with a Holm-Sidak’s multiple comparisons test. To examine mutant survival, Mendelian ratio was confirmed for both larvae and adult offspring from a heterozygous incross by Chi-squared tests. For all other quantitative experiments, data were pooled from the independent replicates and results were summarized in terms of least-squared adjusted means (lsmeans) and standard errors (Vincent et al., 2016). Results were analyzed using ANOVA with a Tukey’s multiple comparisons test. Graphical representation shows individual data points color coded to reflect replicates. Statistical analysis and graphical representations were done using R version 3.4 and GraphPad Prism version 6.

### Summary of supplemental material

Supplemental material includes additional data characterizing the zebrafish Padi2 transcripts and proteins (Fig. S1), additional characterization of the *padi2* mutant (Fig S2), and additional non-wound phenotypes observed in the mutant and supporting morpholino data (Fig. S3). We also include 6 videos that characterize the 3 dimensional context of the citrullinated histones post injury.

## Supplemental material

**Figure S1: Characterization of zebrafish Padi2.** (A) Schematics of *padi2* transcripts, with exons represented by solid boxes and introns by connected lines (slashes indicate shortening of relative length for display purposes). Left, list of the corresponding last 7 digits of Ensembl ID from GRCz11 and GRCz10 genome assemblies (full Ensemble IDs listed in materials and methods). Right, list of the names based on GRCz10 used to reference the transcripts. Cloned transcripts discussed in this paper are in green and arrows highlight exon 10. (B) Full amino acid sequences of human PAD2 and predicted zebrafish Padi2 splice variants (201a and 202). Amino acids are highlighted (as indicated in key) to demonstrate calcium binding, catalytic residues, and substrate-binding residues. Black arrow heads indicate amino acids referred to in Fig 1 B and S1 F. (C) Full western blot (from Fig 1 F) of pooled larvae probed with antibodies against zebrafish Padi2 and Actin. Wild-type and *padi2*^−/−^ lysates were probed. Arrow demonstrates expected size of Padi2 transcripts at ∼75 and 80 kDa and asterisk marks ∼200 kDa species. Representative blot from 4 replicates. (D) Western blot of pooled wild-type larvae probed with pre-immune serum and Actin antibody. (E) zPadi2 western blot of pooled 2 dpf larvae. Lane 1, wild-type; lane 2, *padi2^−/−^*; lane 3, *padi2* 201a mRNA-injected *padi2^−/−^* larvae. (F) Citrullination activity of Padi2 202 and individual point mutations in select calcium-binding and catalytic amino acids (colors correspond to highlighted residues in B). Fold change of enzymatic activity normalized to wild-type Padi2 202. Data represent 2 independent experiments and wild-type values are also represented in Fig 1 A.

**Figure S2: Homozygous *padi2* mutants are viable and have increased neuromuscular junctions.** (A) Genotype frequency at 5 dpf larvae of incrossed *padi2* heterozygotes. (B) Genotype frequency of adult offspring of incrossed *padi2* heterozygotes. Data in A and B are from four and six clutches, respectively, and analyzed by Chi-squared tests. (C) Representative images of slow-muscle fibers immunostained with α-MyHC antibody in the trunk of 5 dpf larvae from 3 independent replicates. (D) Representative images of the trunk of phalloidin-stained 5 dpf larvae for visualization of F-actin in fast-muscle fibers. Wild-type cousin (left) and *padi2*^−/−^ (right) from 3 independent replicates. (E) Neuromuscular junctions are labeled with α-SV2 (green, presynaptic vesicles), α-BTX (red, postsynaptic AChRs), and merge (synapses) in wild-type cousins (top) and *padi2*^−/−^ (bottom) larvae at 5 dpf. (F) Quantification of the number of SV2 puncta, AChR puncta, and synapses in a single myotome in the trunks of larvae. Data are from three pooled independent replicates with the lsmeans (±) SEM and p values calculated by ANOVA reported. Each symbol represents a single myotome and measurements were taken from two myotomes per larva (n = 100 myotomes from 50 wild-type larvae, n = 114 myotomes from 57 *padi2*^−/−^ larvae). Scale bars = 50 μm.

**Figure S3: Padi2-deficient larvae show regeneration defects.** (A) RT-qPCR of *padi2* exon5/6 on pooled fin extracts from 24 hpw and no wound controls (3dpf) normalized to wildtype, no wound fins. Data are from three pooled independent replicates with the means and SEM reported and a one-sample t test performed. (B) Quantification of regenerative and developmental fin length after morpholino (MO) knockdown of *padi2*. Data from 5 independent replicates with 3 dpw (n = 90 control MO, n = 113 *padi2* MO) and 5 dpf (n = 104 control MO, n = 103 *padi2* MO). (C) Quantification of neutrophils in whole larvae from 3 independent replicates, (n = 30 +/+, 29 -/- at 2 dpf; n = 30 +/+, 30 -/- at 3 dpf). (D) Quantification of neutrophil numbers in developmental, unwounded fins. Pooled from five independent replicates (n = 88 +/+, 78 -/- at 2 dpf and n = 79 +/+, 75 -/- at 3 dpf). (E) Quantification of macrophage numbers in whole larvae from 3 independent replicates (n = 30 +/+, 29 -/- at 2dpf and n = 30 +/+, 30 -/- at 3 dpf). (F) Quantification of macrophage numbers in developmental, unwounded fins. Pooled from 4 independent replicates (n = 81 +/+, 74 -/- at 2dpf and n = 70 +/+, 66 -/-). (G) Representative images of H4cit3 immunostaining in 3 dpf unwounded control larvae with H4cit3 antibody label on the right and merged with the bright-field on the left. (H) Representative multiphoton microscopy 3D reconstruction showing enface view of the notochord bead at 24 hpw in *Tg(lepb:EGFP)* expressing (green) larvae labeled with H4cit3 immunofluorescence (magenta). Last image in row includes brightfield overlay. (I) Section view of the notochord bead, showing enface (x,y view) and orthogonal (x, z view is below; y, z view is to the right) sections, with section thickness shown 2 μm for both x and y, 10 μm in z. Scale bars = 10 μm. (J,L) Representative images of active-Caspase3 labeled in (J) 66hpw fins or (L) developmental, unwounded fins. Merged images of active-Caspase3 (magenta) and DAPI (white) on the left, and single active-Caspase3 channel in white on the right. (K) Quantification of active-Caspase3 threshold area in *padi2^−/−^* and wild-type fins at 66 hpw from 3 independent replicates (n = 47 +/+, 47 -/-). (M) Representative images at 24 hpw and (N) quantification of mitotic cells labeled with phosphorylated histone H3 in MO injected larvae past the notochord (white dotted line) from 3 independent replicates (24 hpw n = 68 control MO, 70 *padi2* MO and 3 dpf n = 71 control MO, 64 *padi2* MO). All quantifications have lsmeans and SEM reported and p values were calculated by ANOVA. Scale bars = 100 μm.

## Acknowledgements

We thank members of the Huttenlocher lab for helpful discussions of the research as well as with technical support and zebrafish maintenance. We thank Dr. Emily Rosowski for her careful reading of the manuscript and suggestions and Dr. Laurel Hind for her critical edits of the manuscript. We would like to thank Francisco Barros Becker for assistance with FIJI analysis and Jens Eickhoff for advice on statistical analyses.

This work was supported by NIH R35 GM1 18027 01 to AH. NG was supported by a Molecular Biosciences Training Grant T32-GM07215, JK was supported by the American Heart Association grant (AHA16SDG30020001) and MAS by NIH/NIAMS K08 AR065500. The authors declare no competing financial interests.

## Author contributions

NG, JMS, MAS, KWE, JK and AH conceived and designed experiments. NG, DAB, JMS, JR, and PP conducted the experiments. NG and DAB performed the analysis. NG, JMS, and AH prepared the figures and wrote the manuscript.

