## Supplemental figures for "Citrullination regulates wound responses and tissue regeneration in zebrafish"

### Figure S1: Characterization of zebrafish Padi2

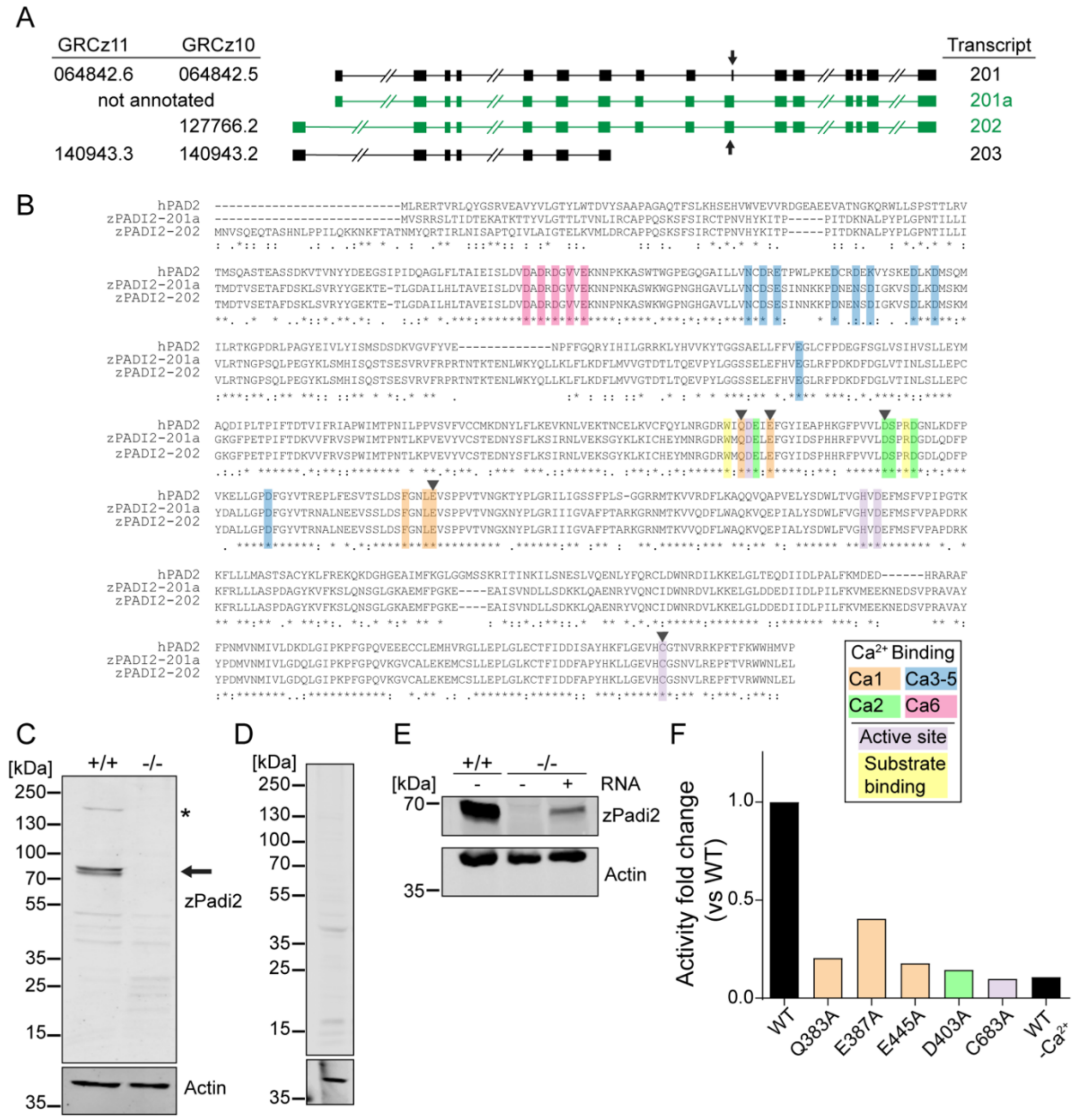

**Figure S2: Homozygous *padi2* mutants are viable and have increased neuromuscular junctions**

A

| Genotype | **+/+** | **+/-** | **-/-** |
|---|---|---|---|
| Total number | 298 | 609 | 290 |
| Percentage | 25% | 51% | 24% |

B

| Genotype | **+/+** | **+/-** | **-/-** |
|---|---|---|---|
| Total number | 41 | 56 | 29 |
| Percentage | 33% | 44% | 23% |

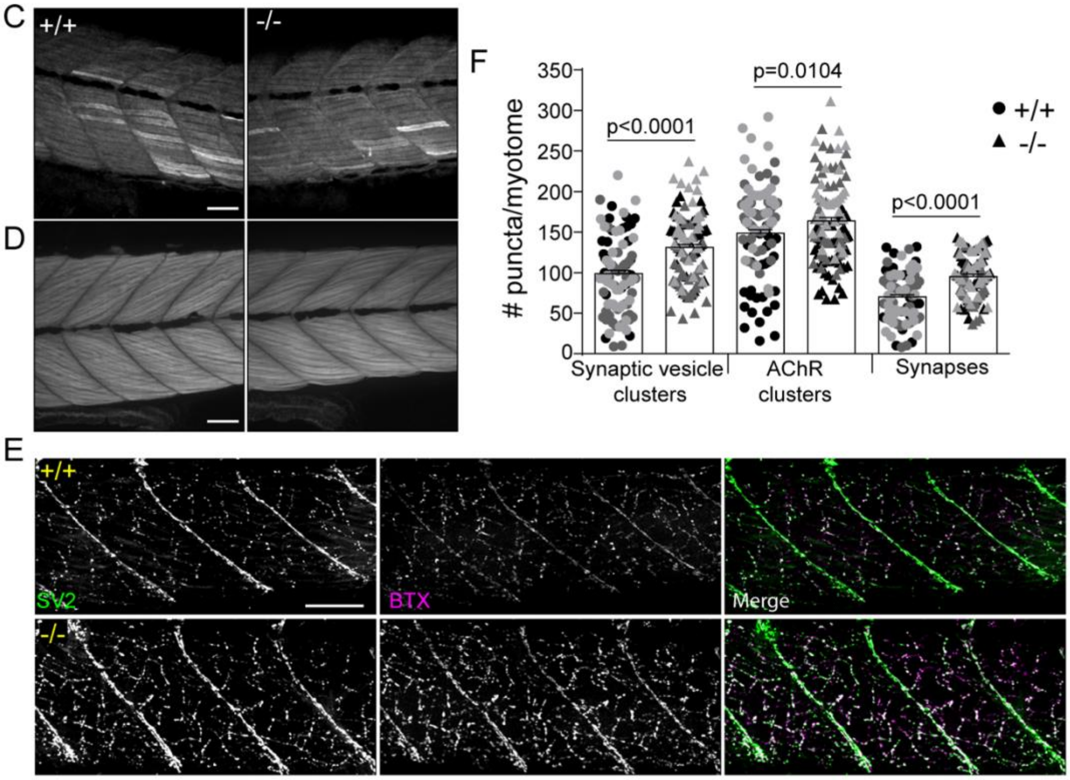

**Figure S3: Padi2-deficient larvae show regeneration defects**

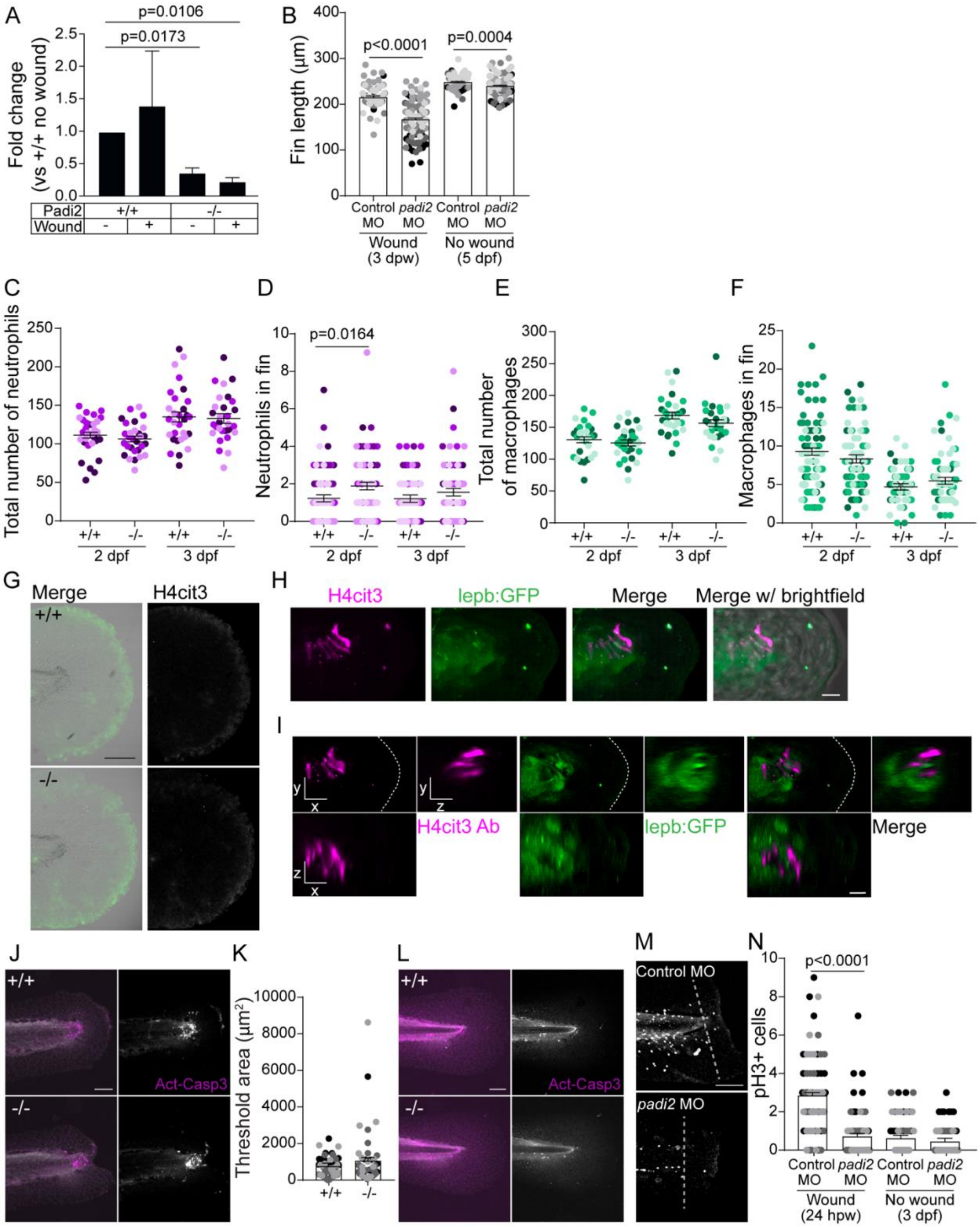
